# Bagworm decorations are an anti-predatory structure

**DOI:** 10.1101/2019.12.12.873893

**Authors:** Md Kawsar Khan

## Abstract

Many animals decorate their exterior with environmental materials and these decorations are predicted to increase their survival. The adaptive significance of these decorations, however, has seldomly been tested experimentally under field conditions. Here, I studied the anti-predatory functions of the decoration (bag) of a bagworm moth, *Eumeta crameri* against their natural predator, *Oecophylla smaragdina*, the Asian weaver ant. I experimentally tested if bag removal from caterpillars resulted in more predation than bagged caterpillar under field conditions, which would support the hypothesis that bags are selected to protect the caterpillars against their predators. In support of that, I showed that caterpillars without a bag were attacked, killed and taken to ants’ nest significantly more than bagged caterpillars. My study provides rare experimental evidence for anti-predatory functions of the decoration. My study suggests that decorating behaviour has evolved in animals as an anti-predatory defence mechanism.

## Introduction

Many animals actively accumulate and attach environmental materials to their exterior (Ruxton & Stevens, 2015). For example, rodents apply the shed skin of snakes to their fur, crabs carry coconut shells, assassin bugs carry carcass of their ant prey, and caddisfly decorate their cases by attaching debris (Brandt & Mahsberg, 2002; Clucas *et al.*, 2008; Hultgren & Stachowicz, 2008; Ferry *et al.*, 2013). These decorations are predominantly considered to have a defensive function against predators; however, their adaptive functions have rarely been tested directly, especially, under field conditions (Ruxton & Stevens, 2015).

Lepidopteran larvae of the Psychidae family (the bagworm moths) construct bags with silk and decorate them with environmental material such as leaves, grasses, bark fragments, wood debris, twigs, lichens, and sand particles (Rhainds, Davis & Price, 2008; Sugimoto, 2009). Bagworm caterpillars carry these bags while foraging. The bags generate a self-enclosing microclimatic condition (such as increased temperature) that increase overwinter survival, accelerate development and reduce desiccation (Smith & Barrows, 1991; Rivers, Antonelli & Yoder, 2002; Rhainds *et al.*, 2008). Furthermore, bags are believed to function as a physical barrier to protect the caterpillars from parasitoid and predator attack. There is evidence for that bags reduce parasitoid attacks, such as, large bags of *Thyridopteryx ephemeraeformis* were shown to reduce parasitoid attacks (Cronin & Gill, 1989). Alternatively, Bagworm caterpillars were reported to experience greater parasitism than externally feeding caterpillars without bags (Hawkins, 2005). In addition to parasitoids, birds and ants predate on the bagworm moths (Moore & Hanks, 2000; Pierre & Idris, 2013). Which suggests that the bags may have evolved to protect the bagworm moths from predators. In support of this idea, bags of *Eumeta minuscula* moths provided protection against their putative predator *Calosoma maximoviczi* under laboratory conditions (Sugiura, 2016). However, anti-predatory functions of the bags have not been yet studied against their natural predators under field conditions.

Here, I studied the potential anti-predatory function of bagworm bags using *Eumeta crameri* as a model system. I removed the bags from the caterpillars and placed the caterpillars on the naturally occurring trees with bagged-caterpillars and determined the predation rate from their natural predators *Oecophylla smaragdina.* I predicted caterpillars without their bags will be predated more than the bagged caterpillars, if bags function to protect against predators.

## Materials and methods

### Study species

*Eumeta crameri* is a widely distributed moth species in south Asia. In Bangladesh, this species is known to occur in the central region (Dhaka) and its surrounding areas (Ameen & Sultana, 1977). In this region, *Eumeta crameri* has four generations per year, where the caterpillars appear in April, July, October, and December (Ameen & Sultana, 1977). The population of this species is at maximum density between April to July (Ameen & Sultana, 1977). *Eumeta crameri* caterpillars feed on leaves of several woody plants and shrubs (Farooqui & Singh, 1975; Ameen & Sultana, 1977). The newly emerged caterpillars make their bag within 8-12 hours (Ameen & Sultana, 1977). The bags are usually made from caterpillar silk covered by plant materials such as leaves, twigs and bark (Ameen & Sultana, 1977; Thangavelu & Ravindranath, 1985). The caterpillars carry their bags when they search for food plants (Ameen & Sultana, 1977). The moving caterpillars body remains hidden inside the bag with only the head and prothorax visible. The caterpillars are assumed to renovate the bags as they grow. *Eumeta crameri* caterpillars are parasitized by parasitic wasp of the *Brachymeria* genus (Ameen & Sultana, 1977).

### Study sites

I collected *Eumeta crameri* caterpillars in bags from an area of approximately 500 square meters at Sultanpur, Palash, Narsingdi, Dhaka (24°0′N, 90°39′E, 40 meters above sea level). I examined the stems of the plants from the ground up to 2.5 meters to collect the bags. Collections took place between May 27 to June 30 when the abundance of this species was maximal. The bags were collected from a mean height of 112.57 ± 7.74 cm from ground (n = 64) and the average width of the stems were 87.67 ± 3.88 cm (n = 64).

I did not required permission to collect *Eumeta crameri* bags as this species is neither endangered nor protected in Bangladesh and the study area was not part of any national park or protected area.

### Bag-caterpillar correlation and bag renovation experiment

I aimed to determine if bag length and width correlated with caterpillar length and width. I measured the length and maximum width of the collected bags (n =21) using a slide calliper. Then, I removed the bags and measured the length and width of the caterpillars (n =19). A positive correlation between bag size and caterpillar size suggest that caterpillars enlarge their bags as they grow. Further, I experimentally tested whether caterpillars can rebuild their bags. I removed the bags and reared the caterpillars in a rearing box with bark and twig fragments of their host plants. I kept a soaked cotton ball inside the rearing box for moisture and drilled holes into the lid to maintain air flow. I kept the caterpillars under ambient temperature (25-32°C) overnight and observed the following morning if they rebuilt the bags.

### Antipredation experiment

During the collection of *Eumeta crameri* from their host plants, I observed *Oecophylla smaragdina* (Asian Weaver ant), and their nests on those plants. I observed that weaver ants fed on small insects such as dragonflies, butterflies, moths and lepidopteran larva. Previous studies have shown that *Oecophylla smaragdina* are natural predators of *Eumeta crameri* caterpillars (Pierre & Idris, 2013). I aimed to determine if the bags of *Eumeta crameri* caterpillars provide protection from the weaver ants and accordingly predicted that bagged-caterpillars will be predated less than caterpillars without bags.

I removed the bags from the caterpillars and placed the caterpillars 25cm above or below a randomly selected naturally occurring bagged caterpillar on a tree. In this setting, both the caterpillars with and without a bag could move freely on the tree that also housed naturally occurring weaver ants. I observed the caterpillars for 40 minutes and recorded their interaction with the ants. I counted ant predatory responses (attack or no attack) when legs, antenna or any parts of the ants touched the caterpillars. If an ant bit the caterpillar, I counted it as an attack. On the other hand, if the ant left the caterpillar without biting, I counted it as a no-attack. After an attack, if the ant killed and took the caterpillars to their nest, I counted it as successful predation. Otherwise, if the ant failed to kill the caterpillar, I counted it as failed predation. I counted the number of attacks and predation on the caterpillars with and without bag. I performed 25 trials in total, each time with a new caterpillar. I haphazardly selected the trees, and position (above or below) of the naturally occurring caterpillars with bags to introduce caterpillars without bags.

### Statistical analyses

I applied linear models (LMs) to determine the relationship between bag length and caterpillar length, and bag width and caterpillar width. I applied generalized linear mixed effect model (GLMM) to determine whether caterpillars without bags were likely to incur more attacks than caterpillars with bags. I fitted GLMMs with *attacks* as response variables, caterpillar types (bag removed or present) as a fixed factor, and trial identity as a random effect. I used the r.squared GLMM function of the R package ‘MuMIn’ to determine the effect size of the models (Johnson, 2014; Bartoń, 2019). To account for zero inflation and overdispersion, I fitted zero-inflated generalized linear mixed model (ZIGLMM) to determine if caterpillars without bags were predated more than those with a bag. I fitted ZIGLMMs with predation as response variables, caterpillar types (bag removed or bagged) as a fixed factor, and trial identity as a random effect. I performed all analyses in R v 3.5.2 using packages ‘lme4’ (Bates *et al.*, 2019), ‘glmmADMB’ (Bolker *et al.*, 2012), ‘yarrr’ (Phillips, 2017) and ‘MuMIn’(Bartoń, 2019).

## Results

### Bag-caterpillar correlation and bag renovation

Caterpillar length was significantly correlated with the length of the bag (LM: estimate = 2.13 ± 0.15, *t* = 13.84, *p* < 0.0001, R^2^ = 0.90; Figure 1c). Similarly, caterpillar width was correlated with the bag width (LM: estimate = 2.98 ± 0.22, *t* = 4.44, *p* < 0.001, R^2^ = 0.58; Figure 1d). Following bag removal, nine out of ten caterpillars rebuilt their bag overnight with the plant fragments.

**Figure 1:**
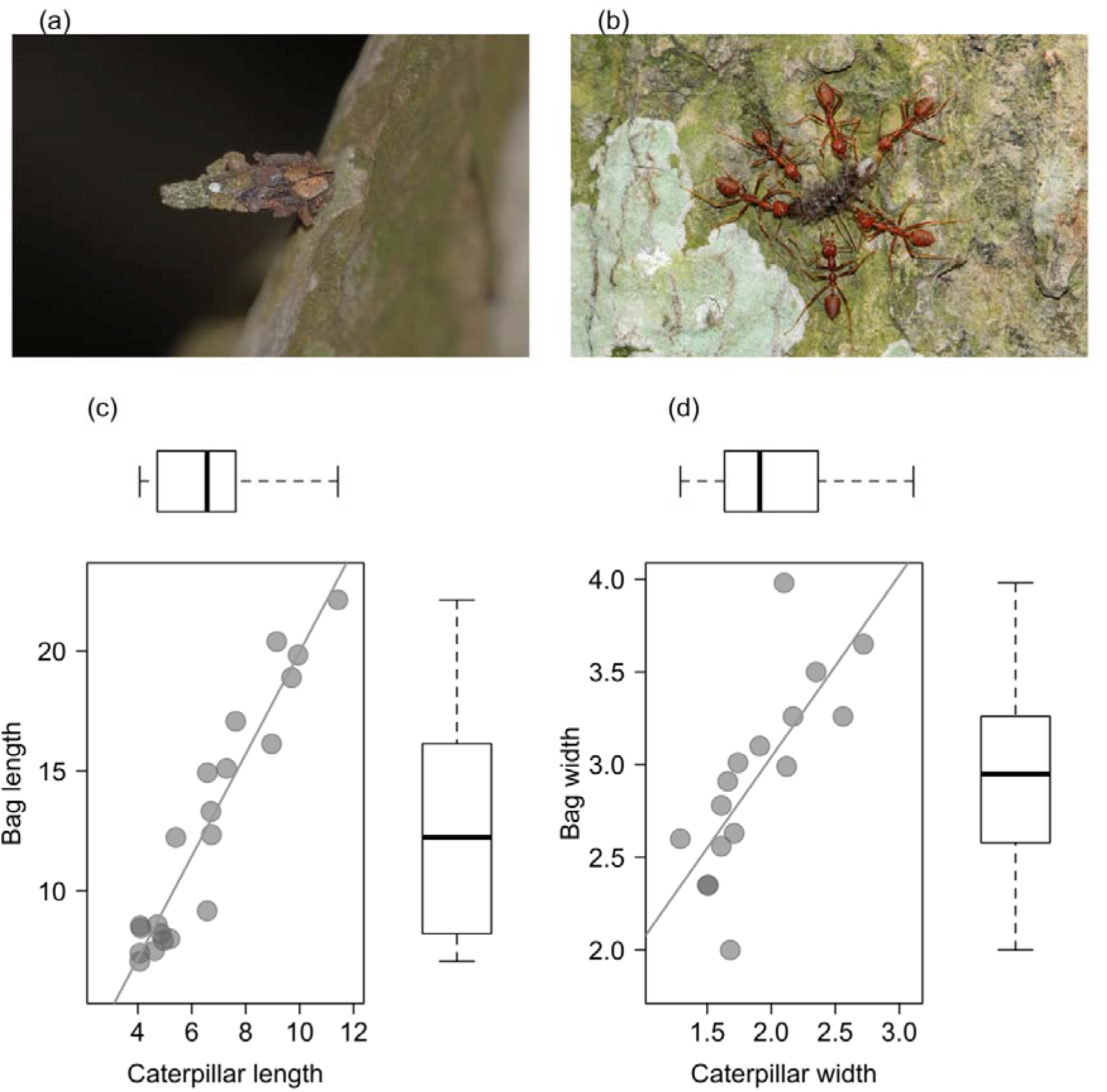
a) Photograph of a *Eumeta crameri* bagworm, b) photograph of *Oecophylla smaragdina* predating a caterpillar, c) scatterplot showing the correlation between the bag length and caterpillar length; upper boxplot shows the distribution of bag length (n = 21) and right boxplot exhibits the distribution of caterpillar length (n = 21), d) scatterplot showing the correlation between the bag width and caterpillar width, upper boxplot exhibits the distribution of bag width (n = 19) and right boxplot exhibits the distribution of caterpillar width (n = 19). Solid line represents the regression line. Bold lines of boxplot indicate medians; boxes enclose 25th to 75th percentiles. The whiskers extend to the data range, excluding outliers beyond 1.5 times the interquartile range.

### Predation experiment

The bagged caterpillars were attacked less frequently than the caterpillars without bag (GLMM: estimate = −14.39 ± 5.45, *z* = −2.63, *p* < 0.01, R^2^ = 0.58; Figure 2a). Similarly, caterpillars with bags were predated (killed and taken to nest) significantly less frequently than those without a bag (ZIGLMM: estimate = −2.78 ± 1.00, *z* = −2.77, *p* < 0.01; Figure 2b).

**Figure 2:**
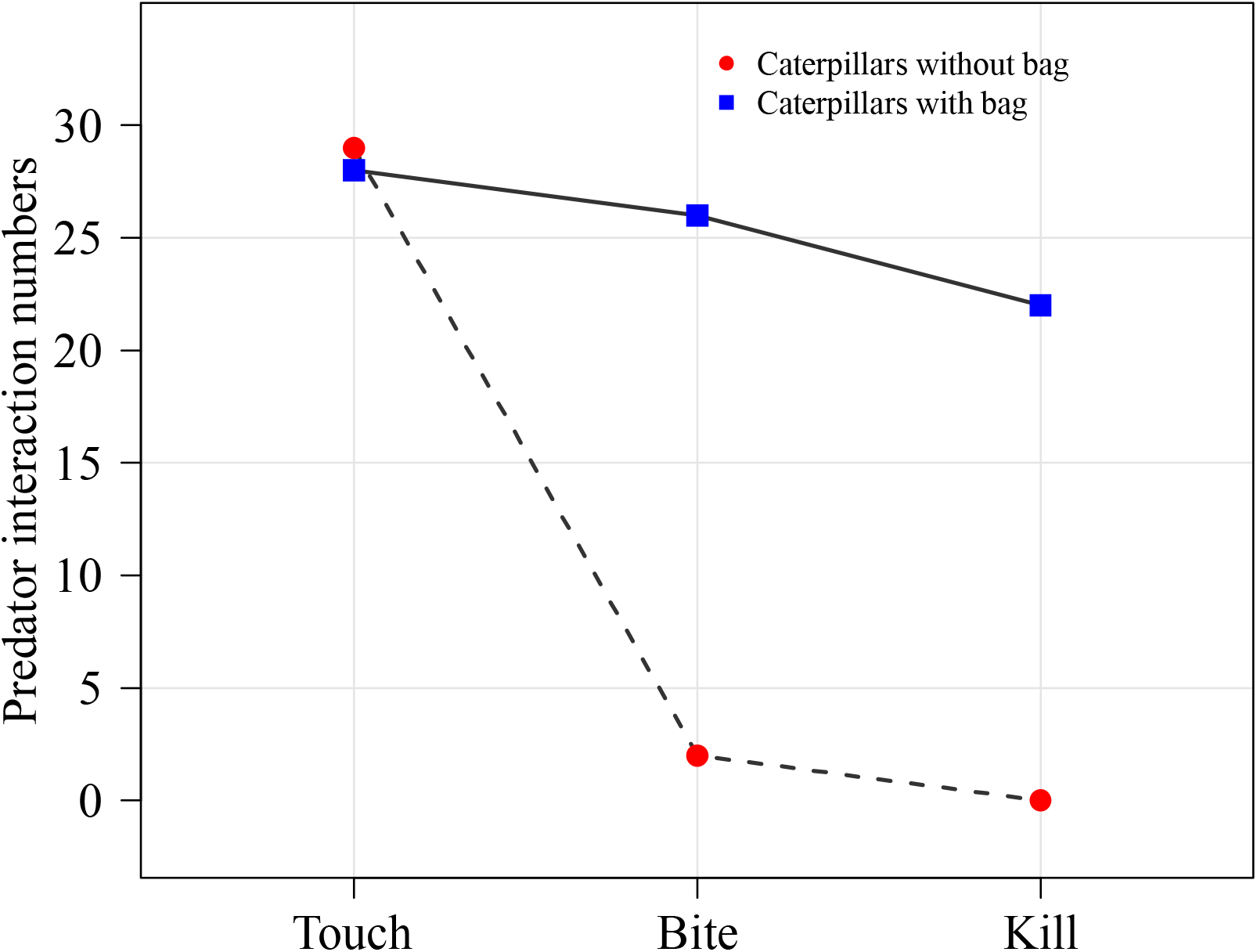
Numbers of predator interactions (touch, bite and kill) with the bagged and bag removed caterpillars (n =25 trials).

## Discussion

Animal decorations such as the bags of the bagworm moths are predicted to provide protection from predators, however, their function has rarely been tested experimentally under field conditions. Here, I showed that the bags of the *Eumeta crameri* moths provide protection against their natural predators *Oecophylla smaragdina*, the Asian weaver ant. I experimentally showed that caterpillars without bags were attacked and predated more than those with a bag.

Animal decorations often conceal the decorators via background matching, masquerade, and disruption (Ruxton & Stevens, 2015). My data show that caterpillars without bags were equally likely to be touched by ants as the bags, suggesting that ants were able to detect both (Figure 2). However, the weaver ants, in my experiments, did not attacked the bagged caterpillars even after touching the bags. Possibly, the ants did not detect the caterpillars within the bag suggesting that the bags reduced the detection of the caterpillars as potential prey. Alternatively, ants may have detected the caterpillar within the bag, but decided not to attack. It is likely that ants failed to detect the caterpillars in the bag as on several occasions; after touching the bags, the ants left without biting the caterpillars. The bags probably masked cues of the caterpillar by masquerade and the ants misclassified the bag as part of a plant. Furthermore, the chemical compositions of the bags possibly also masked chemical cues of the caterpillars. However, further studies are required to understand the precise mechanism of recognition.

Even when a predator recognises prey cue, decorations can function as physical barriers to protect the decorators (Ruxton & Stevens, 2015). My study showed that weaver ants could not break the bags with repeated biting, even when they detected the bags as potential prey. However, the ants easily killed caterpillars without a bag and took them to their nest. These experimental observations, affirms that the bags function as a protective barrier against ants’ predation. Similar protection via physical barriers were shown in Antarctic sea urchins; hydroid decorated sea urchins survived anemone attack but urchins without hydroids were killed in a repeat encounter (Dayton, Robillard & Paine, 1970). Even if ants could break through the physical barrier, bags can still deter predators because of the longer handling time required to remove the bags. For example birds prefers caterpillars without bags over those with bags to avoid greater processing cost (Moore & Hanks, 2000).

Decorating is an intriguing behaviour that has evolved in many taxa. Bag decorating behaviour occurs in more than 1000 species of family Psychidae (Lepidoptera: Tineoidea) (Rhainds *et al.*, 2008). The bag decorations are often assumed to provide protection against predators, however, their adaptive significance has been rarely tested experimentally under natural conditions. Here, I have shown that the bags of in *Eumeta crameri* provide protection against their natural predators under field conditions. My study provides evidence for the anti-predatory function of the bags. The bags, however, could have other potential functions such as temperature regulation, protection from dehydration, and parasitoid attacks. Future studies should test these mutually exclusive functions together to determine if their synergistic functions reinforce the evolution of bag decorations.

## Acknowledgements

I thank Marie Herberstein for commenting on the initial version of the manuscript. I thank Payal Barua for all her support.

## Data accessibility

All data will be accessible upon publication

